# High molecular-weight polysaccharide contamination from yeast extract in semi-defined bacteriological media: Effects on exopolysaccharide production and purity

**DOI:** 10.64898/2026.02.27.708493

**Authors:** Ahmad Tsjokajev, Gordon Jacob Boehlich, Svein Jarle Horn, Gustav Vaaje-Kolstad, Bjørge Westereng

**Affiliations:** Faculty of Chemistry, Biotechnology and Food Science, Norwegian University of Life Sciences (NMBU), Ås, Norway

**Keywords:** Bacterial exopolysaccharides (EPS), Lactic acid bacteria (LAB), MRS medium, Yeast extract, Yeast mannan

## Abstract

Exopolysaccharides (EPS) produced by lactic acid bacteria (LAB) and other microorganisms have attracted considerable interest due to their structural diversity and physicochemical properties, which makes them valuable across various industrial applications. To achieve high cell densities and maximize EPS yields, microorganisms are typically cultivated in nutrient-rich media containing yeast extract. However, yeast extract may contain high molecular weight polysaccharides that are not metabolized by the bacteria. This can lead to an overestimation of EPS yields and contamination of the bacterial EPS, potentially resulting in misinterpretation of their structure and biological activity. In this study, we demonstrate the presence of high molecular weight α-mannan and β-glucan in yeast extract in EPS isolates using both ultrafiltration and the commonly used trichloroacetic acid/ethanol (TCA/EtOH) precipitation method. These polysaccharides were characterized by size-exclusion chromatography, high-performance anion-exchange chromatography, and nuclear magnetic resonance spectroscopy. Their abundances were estimated to range from 10 to 50 mg/L in MRS medium, depending on the supplier of the yeast extract. The main contaminant identified was yeast α-mannan. By cultivating *L. rhamnosus* GG (ATCC 53103) and *L. pentosus* KW1 and isolating their respective EPS, we illustrate how these yeast extract contaminants affect the structural interpretation of the EPS and that the contaminants can be completely removed by ultrafiltration of the growth medium prior to bacterial cultivation. In conclusion, we emphasize the necessity of stringent controls in the production and purification of microbial EPS, with particular attention to the chemical purity of medium constituents.

## 1. Introduction

Exopolysaccharides (EPS) produced by bacteria have current and potential applications in the food, pharmaceutical, cosmetic, and biotechnology sectors [4,33]. For example, EPS produced by lactic acid bacteria (LAB) during *in situ* fermentation play important roles as natural bio-thickeners, gelling agents, stabilizers, water-binding agents and viscosity enhancers in food products. These properties make them particularly relevant in dairy products, as well as in the pulses (e.g. soy), cereal, and baking industries [10,11,22,28]. Further, EPS have been associated with potential health-promoting properties, including antimicrobial, and immunomodulatory activities [10,14,22,33].

LAB are fastidious microorganisms that require complex and nutrient-rich media for optimal growth. Due to the diversity among LAB, no universal culture medium exists. However, one of the most widely used media for cultivating LAB is De Man, Rogosa and Sharpe (MRS) medium, along with modified variants such as M17 [17]. MRS is a semi-defined medium containing specific salts, an emulsifier, nitrogen and carbon sources. In MRS, as well as in many other commonly used media for general bacterial cultivation, yeast extract is frequently used as a source of amino acids, peptides, nucleic acid derivatives, minerals, and vitamins [17].

Yeast extract, an autolysate of *Saccharomyces cerevisiae*, has been reported to contain α-mannan and β-glucan, both of which are constituents of the yeast cell wall [2,7,45]. In the context of EPS production, such polysaccharides can lead to contamination and overestimation of EPS yields if they are not metabolized by the bacteria and instead accumulate in the fermentation broth [7,43].

Uninoculated MRS broth containing yeast extract, beef extract, and proteose peptone has been shown to have a total polysaccharide content of ∼500 mg/L, with yeast mannan identified as the predominant contributor [20,38]. Furthermore, it has been demonstrated that mannans found in EPS fractions of six cultivated basidiomycetes may originate from yeast extract in the growth medium, rather than being synthesized by the fungi themselves [21]. Some studies have addressed the contamination challenge by using ultrafiltration (UF) to remove yeast mannan from the yeast extract prior to adding the filtrate to the cultivation medium for EPS production [5,31]. Unfortunately, these key findings have been largely overlooked, and numerous publications have emerged (**Table 1**), and continue to emerge, that report EPS from a wide range of bacteria containing mannans with structures identical to that of yeast α-mannan [47].

**Table 1.** An overview of studies reporting microbial EPS that may be contaminated by mannan based on the methods and results described. Lactobacillus studies (gray highlight), Gram-positive bacteria (green highlight), Gram-negative bacteria and fungi (blue highlight). Abbreviations: YM, yeast mannan; YE, yeast extract; WP, whey permeate; LB, Luria-Bertani broth; NB, nutrient broth.

| Species/Strain | Medium<sup>1</sup> | Monosaccharide Composition<sup>2</sup> | Mw in kDa | NMR confirmation of Yeast Mannan | Ref. |
| --- | --- | --- | --- | --- | --- |
| *Lactiplantibacillus plantarum* VAL6 | MRS | Man:Glc:Rha:Gal (72:23:3:2) | - | NMR data is not interpretable due to line broadening and low S/N ratio. | [34] |
| *Lactiplantibacillus plantarum* 70810 | YE | EPS1 and EPS2 are composed of Glc:Man:Gal (18:79:3 and 13:31:56) | 205 and 203 | YM is likely present as contaminant in EPS1. | [48] |
| *Lactiplantibacillus plantarum* BC-25 | MRS | Man:Gal:Glc (92:2:6) | 18.3 | No peak assignments. Heavily contaminated with non-carbohydrates, likely YM present. | [53] |
| *Lactiplantibacillus plantarum* HMX2 | MRS | Glc:Gal:Man (43:10:47) | 40 | No NMR data. | [50] |
| *Lactiplantibacillus plantarum* MTCC 9510 | YE | Glc:Man (2:1) | - | <sup>1</sup>H-NMR data not shown, but peak description matches YM. | [19] |
| *Lactiplantibacillus plantarum* W1 | MRS+YE | Glc:Man (3:2) | 111 | NMR suggests presence of YM as contaminant | [12] |
| *Lactobacillus helveticus* MB2-1 | WP+YE | EPS1, EPS2 and EPS3 composed of Gal:Glc:Man at different molar ratios | 200 | No NMR data. | [29] |
| *Lacticaseibacillus paracasei* 2333<br>*Lacticaseibacillus rhamnosus* 1019<br>*Lactobacillus delbrueckii subsp. bulgaricus* 1932 | Modified MRS | Man and Glc (64 % average total) and others: Rha, Fru, Gal, Glc, GlcN, GalN | Four fractions (10–500) | No NMR data. | [15] |
| *Lacticaseibacillus rhamnosus* ACS5 | MRS | Man:Gal:ManNAc (61:19:20) | 107 and 24 | NMR suggests presence of YM as contaminant | [18] |
| *Lactiplantibacillus paracasei* EPS DA-BACS | MRS | Man:Glc as major component (2:1), and others: Fuc, Rha, Gal, GlcN, Ara | 3700 | No NMR data. | [26] |
| *Lactobacillus crispatus* L1 | YE | - | - | NMR matches that of YM. | [13] |
| *Lactobacillus spp.*<sup>3</sup> | MRS | All EPS mainly Man:Glc (ca. 75:25) | 2-3 fract.0.1-10 0 | No NMR data. | [44] |
| *Lactobacilli strains* | MRS | EPS's were composed of Man:Glc (95:5) | - | No NMR data. | [41] |
| *Streptococcus thermophilus* ASCC 1275 | M17 | Fraction 1-3 contain Man, Glc, Gal, Rib: 9:80:11:0, 62:24:14:0, 12:42:24:22 | > 511, 40 and 5 | YM likely present as contaminant in Fraction 2. | [36] |
| *Enterococcus faecium* MS79 | MRS | Ara:Man:Glc (1:2:11) | 830 | No NMR. | [1] |
| *Enterococcus faecalis* | LB | Man:Glc:Fuc (9:2:1) | 260 | NMR suggests presence of YM as contaminant. | [8] |
| *Bifidobacterium breve* H4-2 | MRS | Man:Glc (90:6) | 39.9 | NMR matches that of YM. | [35] |
| *Pediococcus acidilactici* MT41-11 | MRS | Fraction 1-2 contain Man:Glc; (53:1) and (3:54) traces of Rha, Gal, Fuc, Xyl. | 68 and 69 | NMR matches that of YM. | [3] |
| *Bacillus licheniformis* | MRS | Man:Glc (81:7). Constant EPS yield throughout the growth curve. | - | No <sup>1</sup>H-NMR data is shown. The <sup>13</sup>C-NMR data not interpretable low S/N. | [25] |
| *Tetragenococcus halophilus* | MRS | EPS1 composed of Gal:Glc:Man:GlcA (1:6:3:1), and EPS2 of Glc:Man (1:6) | 2613 and 93 | NMR suggests presence of YM as contaminant. | [51] |
| *Rhodopseudomonas palustris* | Salts + YE | Mannan | 47 | NMR matches that of YM. | [52] |
| *Pseudoalteromonas haloplanktis* TAC 125 | YE | Mannose with traces of glucose | - | NMR matches that of YM. | [9] |
| *Psychrobacter arcticus* 273-4 | LB | Man:Glc | 130 | NMR matches that of YM. | [6] |
| *Shigella flexneri* CMCC51574 | LB | EPS 2-1 composed of Man:Glc:GlcN (0.88:0.10:0.02) | 585 | NMR matches that of YM. | [42] |
| *Agaricus sinodeliciosus* var. *Chaidam* | YE | Man:Gal:Rha:Glc:GalN (64:27:2:1:4) with traces of Fuc, Rib, GlcA | 18 | NMR suggests presence of YM as contaminant. | [30] |
| *Pseudomonas aeruginosa* AG01 LC586427 | NB | Man, Gal, Glc, Raffinose, Ara, Xyl (14, 10, 9, 8, 8, 5 mg/g) | - | No NMR data. | [16] |

<sup>1</sup> alfamannan containing ingredients in growth medium used in the study
<sup>2</sup>The various exopolysaccharides mentioned in the literature, found in this table, use various abbreviations, such as EPS 1, 2 etc. The numbering typically follows the chronological elution order based on SEC, with EPS/Fraction 1 representing the fraction with the highest Mw.
<sup>3</sup> *Lactiplantibacillus plantarum* GD2, *Lacticaseibacillus rhamnosus* E9, *Levilactobacillus brevis* LB63, and *Lactobacillus delbrueckii* ssp. *bulgaricus* B3. They present shift values in table, but no supplementary data of spectra etc.

In this study, we demonstrate the presence of high molecular weight (Mw) polysaccharides in yeast extracts from several commercial suppliers and highlight the potential pitfalls these contaminants pose when evaluating EPS purity and structural composition. Furthermore, we explore growth medium adaptation for EPS production in the commercial strain *L. rhamnosus* GG and the in-house strain *L. pentosus* KW1, highlighting the presence of these contaminants and proposing a potential solution to avoid ambiguous structural characterization of EPS.

## 2. Experimental

### 2.1 Preparation of high Mw polysaccharides from yeast extract by ultrafiltration

20 g of <u>Yeast Extract</u> (1.03753, Millipore), <u>Bacto™ Yeast Extract</u> (212750, Thermo Fisher Scientific) and <u>Oxoid™ Yeast Extract</u> (LP0021B, Thermo Fisher Scientific) were individually dissolved in 1 L Milli-Q water and ultrafiltered using a Pellicon® 2 Mini Cassette Holder (Millipore) equipped with a Biomax® 30 kDa Membrane (P2B030V01, Millipore). The retentates were concentrated, followed by diafiltration with Milli-Q water and subsequent lyophilization.

### 2.2 Preparation of high Mw polysaccharides from yeast extract using TCA/EtOH precipitation

10 g of <u>Yeast Extract</u> (1.03753, Millipore) was dissolved in 100 mL Milli-Q water and treated with 50 % TCA to reach a final concentration of 15 %. The solution was incubated overnight at 4°C. The resulting protein precipitate was removed by centrifugation at 4136 × g for 20 min at 4°C (Megastar 1.6R, VWR, Germany). The supernatant was then filtered using a 0.45 µm pore size Steritop filter (10040-450, VWR). Polysaccharides in the filtrate were precipitated by adding cold 96 % EtOH to a final concentration of 70%, followed by overnight incubation at 4°C. The precipitate was collected by centrifugation at 4136 × g for 20 min at 4°C, and the resulting pellet was dried in an oven at 30°C overnight.

### 2.3 Bioreactor fermentations

#### 2.3.1 Bacterial strains

The bacterial strains used for EPS production were the in-house strain *Lactiplantibacillus pentosus* KW1, originally isolated from green olives [49], and the commercially available *Lacticaseibacillus rhamnosus* GG (ATCC 53103) [24].

#### 2.3.2 Growth medium

The MRS medium used for bacterial fermentation contained 10 g/L Peptone (casein-based), 8 g/L Meat Extract, 4 g/L Yeast Extract, 2 g/L K_2_HPO_4_, 5 g/L CH_3_COONa·3H_2_O (Sodium Acetate Trihydrate), 2 g/L HOC(CO_2_NH_4_)(CH_2_CO_2_NH_4_)_2_ or C_6_H_17_N_3_O_7_ (Ammonium Citrate Tribasic), 0.2 g/L MgSO_4_·7 H_2_O and 0.05 g/L MnSO_4_·H_2_O. All ingredients were purchased from Merck Millipore (Burlington, MA, USA).

High Mw polymer contaminants were removed from the medium by ultrafiltration as follows. Ingredients equivalent to 5 L of MRS were dissolved in 1 L Milli-Q water and ultrafiltered using a Biomax® 30 kDa Membrane until the permeate volume reached 5 L. The permeate (referred to as “UF-MRS”) was sterile filtered using 0.22 µm pore size Steritop filter (SCGPS05RE, Millipore) and used as fermentation medium. Prior to inoculation, the medium was adjusted to pH 6 by titration with 2.5 M H_2_SO_4_ and supplemented with ᴅ-glucose to a final concentration of 20 g/L.

#### 2.3.3 Bioreactor cultivation of *L. rhamnosus* and *L. pentosus*

Fermentations were carried out in 1.5 L Minifors 2 bioreactors (Infors HT, Switzerland) using UF-MRS medium at 37°C with continuous stirring at 200 rpm. The pH was maintained at 6.0 by automatic addition of 5 M NaOH. The inoculum was prepared by transferring bacterial cells from a -80°C glycerol stock into 40 mL of UF-MRS medium in a 50 mL Nunc plastic tube using an inoculation loop. This starter culture was incubated statically at 37°C for 6 hours. Subsequently, a 10-fold dilution series was prepared from the starter culture in 50 mL Nunc tubes (final volume: 40 mL), ranging from 10^0^ to 10^8^. The dilution series was incubated statically overnight at 37°C (18 hours). The following day, the culture with an OD_600_ of approximately 6–7 (exponential phase) was used to inoculate 0.8 L of UF-MRS medium.

#### 2.3.4 Extraction, isolation, and purification of exopolysaccharides

EPS was isolated through a multi-step process. After fermentation, the culture was harvested at the end of the exponential growth phase and heated to 100°C for 10 minutes to deactivate any potential depolymerizing enzymes [37]. The cell suspension was then centrifuged at 15,970 × g for 30 minutes at 4°C using a JLA-8.1000 rotor in an Avanti JXN-26 centrifuge (Beckman Coulter). The resulting supernatant was filtered through a 0.22 µm pore size Steritop filter, followed by ultrafiltration and diafiltration with Milli-Q water using a Biomax® 30 kDa Membrane. The final retentate was lyophilized and stored at room temperature until further use.

### 2.4 Structural characterization of polysaccharides

#### 2.4.1 Polymer analysis by size-exclusion chromatography

High Mw polymers were analyzed using size-exclusion chromatography (SEC). The system included two Yarra SEC columns, SEC 4000 and SEC 2000 (300 × 7.8 mm, 3 µm; Phenomenex, Torrance, CA, USA), connected in series to a Dionex UltiMate 3000 HPLC system (Thermo Fisher Scientific, Waltham, MA, USA). The column setup had an exclusion limit of 1500 kDa. Separation was carried out using 0.1 M Na_2_SO_4_ as the mobile phase, with a flow rate of 0.5 mL/min at room temperature. An injection volume of 20 µL was used for samples prepared at a concentration of 2 mg/mL. Detection was performed using ultraviolet (UV) absorbance at 280 nm with a Dionex UltiMate 3000 VWD-3000 detector for protein and peptide components, and a refractive index (RI) detector (RefractoMax 520, ERC) for non-UV absorbing analytes such as polysaccharides. Column performance was routinely validated using pullulan standards ranging from 1.3 to 800 kDa (Agilent Technologies, Mainz, Germany).

Preparative SEC was performed using an Agilent 1260 Infinity Preparative HPLC system (Agilent Technologies) equipped with Yarra SEC-3000 PREP and SEC-2000 PREP columns (300 × 21.2 mm, 5 µm; Phenomenex, Torrance, CA, USA) connected in series. Detection was carried out using dual-wavelength UV detection (260 and 280 nm) and RI detection. The exclusion limit of the column setup was approximately 700 kDa. The mobile phase consisted of 0.08 M ammonium acetate (NH₄Ac), adjusted to pH 4.0, and was delivered at a flow rate of 5 mL/min at room temperature. Injection volumes ranged from 1 to 4 mL, corresponding to a total sample load of 2–8 mg dry matter. Collected fractions were lyophilized to remove water and NH_4_Ac.

#### 2.4.2 Monosaccharide composition analysis

Monosaccharide composition was performed using high-performance anion-exchange chromatography with pulsed amperometric detection (HPAEC-PAD). Each polysaccharide (see Sections 2.1 and 2.2) at a concentration of 1 mg/mL was hydrolyzed with 1 M trifluoroacetic acid (TFA) at 100 °C for 1 hour. The hydrolysates were dried using a Concentrator plus (Eppendorf, Hamburg, Germany) and reconstituted in Milli-Q water to a final concentration of 1 mg/mL. Samples were further diluted to 0.5 and 0.1 mg/mL prior to analysis. Chromatographic separation was carried out using a Dionex™ ICS-6000 System (Thermo Fisher Scientific, Waltham, MA, USA) equipped with a Dionex™ CarboPac™ PA1 column (2 × 250 mm). The mobile phase consisted of 12 mM KOH, delivered at a flow rate of 0.6 mL/min at 30 °C. An injection volume of 0.4 µL was used. Analytical standards including glucose (Glc), galactose (Gal), glucosamine (GlcN), galactosamine (GalN), *N*-acetyl-glucosamine (GlcNAc), *N*-acetyl-galactosamine (GalNAc), rhamnose (Rha), fucose (Fuc) and mannose (Man), were purchased from Sigma-Aldrich (Merck, Darmstadt, Germany) and prepared using the same procedure as the samples.

#### 2.4.3 Nuclear Magnetic Resonance Spectroscopy

All materials analyzed by nuclear magnetic resonance (NMR) spectroscopy were dissolved in D_2_O and lyophilized to minimize the HDO signal. Then 5 mg of material was dissolved in 0.5 mL D_2_O containing 0.05 wt. % 3 (trimethylsilyl)propionic-2,2,3,3-*d*_4_ acid, sodium salt. ^1^H-NMR and spectra were recorded at 298 K using a Bruker AVANCE III HD 400 MHz instrument equipped with BBO room temperature probe. Chemical shifts are reported in ppm with Na-Trimethylsilylpropanoate-*d*_4_ (δ ^1^H/^13^C = 0.00 ppm) as internal standard.

## 3. Results and Discussion

### 3.1 Commercially available yeast extracts contain high Mw polymers

EPS isolation from culture supernatant is commonly performed by first removing proteins through TCA precipitation, followed by polysaccharide precipitation using either ethanol or isopropanol [27]. However, alcohol precipitation is unspecific and will precipitate all polysaccharides present in the solution, including potential contaminants like mannan, if present [38].

To investigate the presence of high Mw polymers in yeast extract, polymers recovered using both the conventional TCA/EtOH method and UF were analyzed by SEC (**Figure 1**). SEC analysis revealed two distinct peaks corresponding to molecular weights of approximately 50 and 100 kDa, present in varying ratios (**Figure 1A**–**1D**). The negligible UV absorbance associated with these two peaks suggests that these high Mw polymers are polysaccharides.

**Figure 1.**
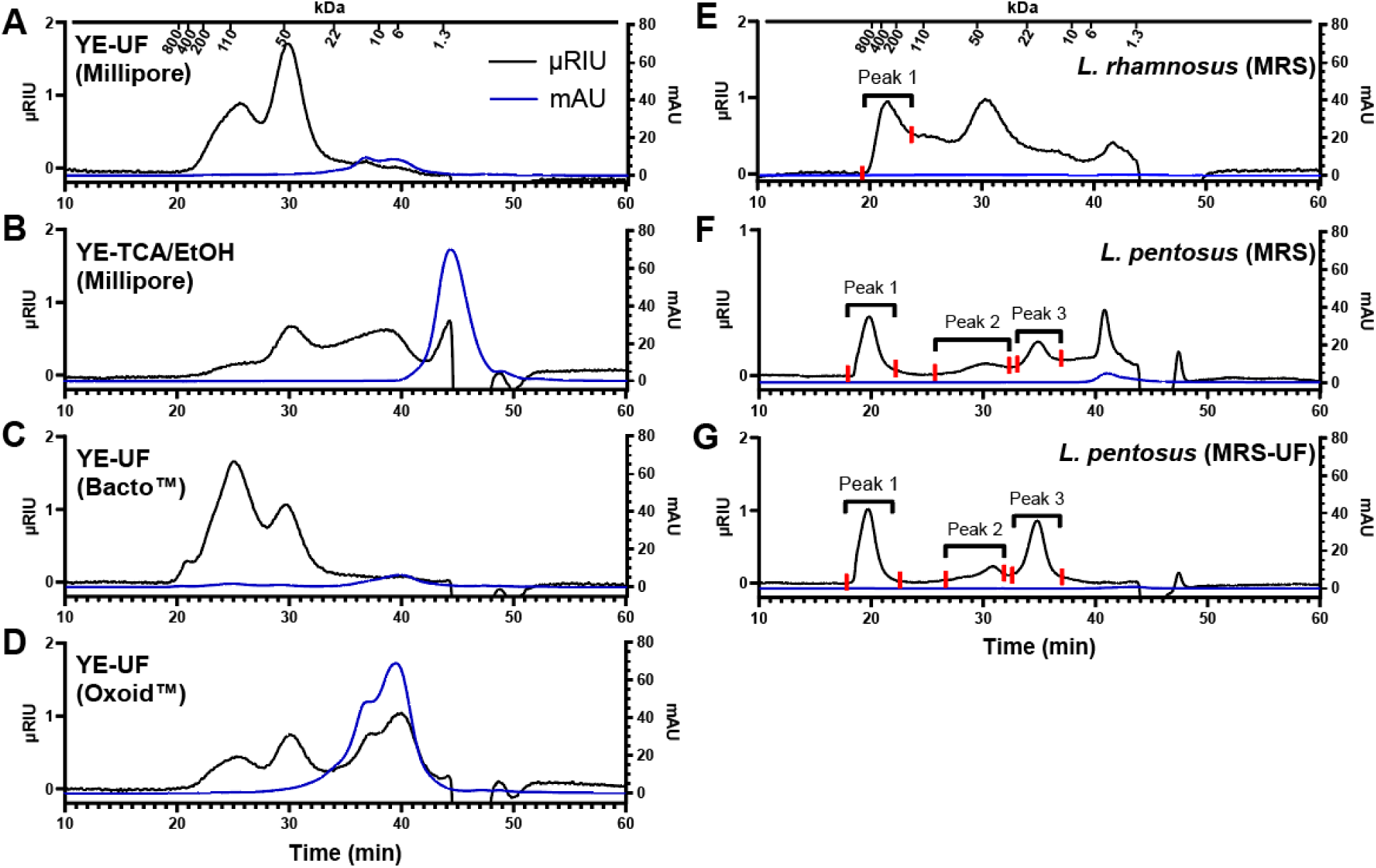
Size-exclusion chromatography (SEC) profiles with refractive index detection (RI, black traces) and ultraviolet detection (mAU_280nm_, blue traces) of recovered polymers from various yeast extracts using ultrafiltration and TCA/EtOH method (**A**–**D**), and of EPS produced by *L. rhamnosus* GG and *L. pentosus* KW1 (**E**–**G**). Red lines in subpanels **E**–**G** indicate the fractionation points used for isolating polymer peaks with preperative SEC (see Section 2.4.1). Separation was performed using a Yarra SEC 4000-2000 series (300 × 7.8 mm, 3 µm) with 0.1 M Na_2_SO_4_ as the mobile phase. YE, yeast extract; UF, ultafiltration; MRS, De Man, Rogosa and Sharpe medium. Peak elution time of pullulan standards (Mw in kDa) are marked above A and E; from left to right 800, 400, 200, 110, 50, 22, 10, 6 and 1.3kDa, respectively.

The SEC profile of yeast extract retentate from Oxoid™ (**Figure 1D**) shows an additional RI signal and a relatively strong UV absorption in the lower Mw range (∼34–43 min), likely due to high Mw peptides retained by the UF membrane. In contrast, the SEC profile of TCA/EtOH-obtained polymers (**Figure 1B**) also shows an additional broad peak (∼33–42 min) with no associated UV-absorbance. This is likely another polysaccharide that is precipitated due to the lower selectivity of ethanol precipitation, which indiscriminately precipitates polysaccharides of all molecular weights. Although SEC analysis does not allow for precise identification of the recovered polymers, the results clearly indicate that yeast extracts contain high Mw polymers, which are co-extracted using both the conventional TCA/EtOH method and the UF method.

### 3.2 High Mw polymers from yeast extract can represent an important contaminant in relevant downstream processed products

To investigate the impact of the high Mw polymers from yeast extract on downstream processes such as EPS production, their carbohydrate content was assessed using data from both SEC-RI and HPAEC-PAD analyses (**Table 2**). For SEC-RI, carbohydrate content was estimated by quantifying peaks corresponding to molecular weight above approximately 22 kDa (retention time < 34 min), based on the assumption that these peaks represent carbohydrates with minimal protein content, as indicated by negligible UV absorbance signals. Carbohydrate content determined by HPAEC-PAD was obtained through quantification of peaks from monosaccharide composition analysis. Carbohydrate content of high Mw polymers from Bacto™ and Millipore yeast extracts was high for both extraction methods (>80 %). In contrast, high Mw polymers from Oxoid™ yeast extract showed a carbohydrate content below 50 %. Similarly, TCA/EtOH-obtained polymers also have reduced carbohydrate content, falling below 50 %. The difference in the carbohydrate content determined by the two analytical methods was small.

**Table 2.** The first section presents the weight of the recovered polymers from three different yeast extracts (YE), obtained using ultrafiltration (UF) and TCA/EtOH method. Carbohydrate purity (%, w/w) and the estimated carbohydrate concentration in MRS medium (mg/L) were determined for each yeast extract using SEC-RI. The second section provides the carbohydrate purity (%, w/w), relative monosaccharide composition (%, w/w), and estimated carbohydrate concentration in MRS medium (mg/L) for each yeast extract, as determined by HPAEC-PAD.

| SEC-RI |  |  |  |  |
| --- | --- | --- | --- | --- |
| Workup/Supplier | Initial ingredient weight (g) | Extracted polymer weight (mg) | Carbohydrate purity<sup>1</sup> (%) | Carbohydrate in MRS medium<sup>3</sup> (mg/L) |
| YE-UF (Millipore) | 20 | 69 | 92 | 13 |
| YE-TCA/EtOH (Millipore) | 10 | 251 | 35 | 35 |
| YE-UF (Bacto™) | 20 | 99 | 98 | 19 |
| YE-UF (Oxoid™) | 20 | 374 | 43 | 32 |
| HPAEC-PAD |  |  |  |  |
| Workup/Supplier | Glc (%) | Man (%) | Carbohydrate purity (%)<sup>2</sup> | Carbohydrate in MRS medium<sup>3</sup> (mg/L) |
| YE-UF (Millipore) | 3 | 97 | 80 | 11 |
| YE-TCA/EtOH (Millipore) | 25 | 75 | 48 | 48 |
| YE-UF (Bacto™) | 17 | 83 | 94 | 19 |
| YE-UF (Oxoid™) | 4 | 96 | 32 | 24 |

<sup>1</sup> Carbohydrate purity of the extracted polymers was determined by quantification of RI-SEC peaks that are above 22 kDa (<34 min), with the assumption that the peaks correspond to carbohydrates with very small amount of protein (negligible UV signal) present. For quantification, pullulan standards (0.1–2 mg/mL) were used for the standard curve.
<sup>2</sup> Carbohydrate purity of the extracted polymers was determined by quantification of monosaccharide peaks using HPAEC-PAD. The calculations were based on absolute mass concentration, while monosaccharide composition was expressed relative to the total carbohydrate content.
<sup>3</sup> The estimated carbohydrate concentration in MRS medium (mg/L), depending on the yeast extract supplier, was calculated based on the extraction yield (derived from the initial ingredient weight and the extracted polymer weight; extraction procedures described in Section 2.1 and 2.2), the carbohydrate purity of the extracted polymers <sup>1,2</sup>, and the amount of yeast extract normally used in the MRS medium (4 g/L).

Based on the calculated carbohydrate content, it was possible to calculate the amount of polysaccharide contributed by the yeast extracts to the MRS medium. Depending on the yeast extract used, a standard MRS medium can have a polysaccharide content ranging from 10 to 50 mg/L (**Table 2**). Assuming that the cultivated microorganism does not consume the polysaccharide present in the MRS medium, this polysaccharide contaminant can represent a substantial part of the EPS obtained from such cultivation, especially from LAB, which typically produce heteropolysaccharides in the range of 50–1000 mg/L [39].

To gain further insight into the identity of the polysaccharides obtained from the yeast extracts, monosaccharide composition analysis (**Figure 2**) revealed a considerable amount of mannose in all polysaccharides (**Table 2**), along with lower amounts of glucose. Additionally, an unidentified peak with a later retention time (∼9.7 min) was consistently observed across all samples, potentially indicating the presence of a modified monosaccharide or a disaccharide resulting from incomplete hydrolysis.

**Figure 2.**
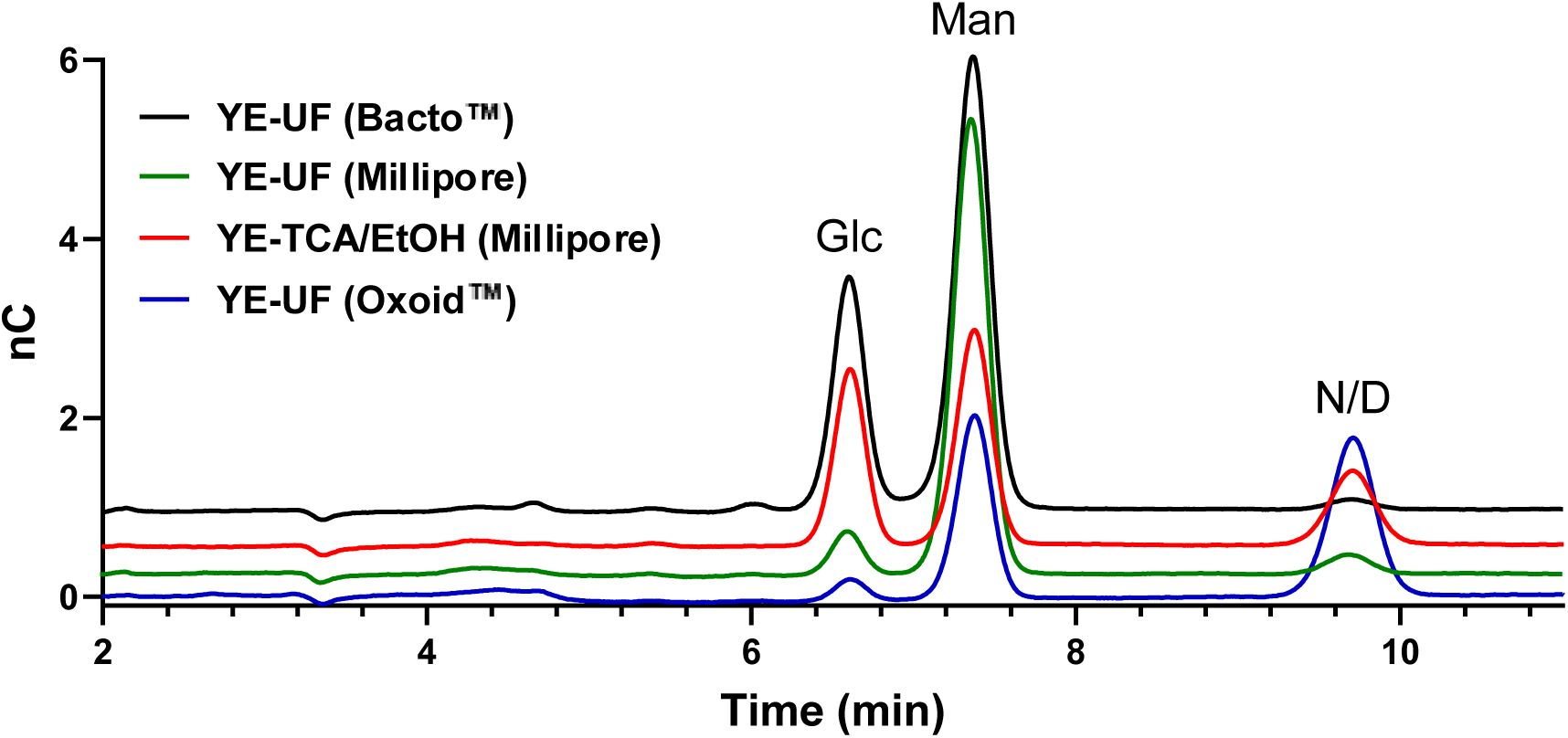
HPAEC-PAD chromatography of polysaccharide hydrolysates isolated from three different yeast extracts (YE) after 1-hour hydrolysis with 1 M TFA at 100 °C. Separation was performed using Dionex™ CarboPac™ PA1 (2 × 250 mm) with 12 mM KOH as the eluent. ND, not determined; Glc, glucose; Man, mannose.

The high mannose content suggests the presence of yeast mannan, while the presence of glucose indicates β-glucans, both of which are structural polysaccharides of the yeast cell wall [32]. The mannose-to-glucose ratios varied slightly for each sample (**Figure 2**), with the lowest ratio observed in the TCA/EtOH extracted polysaccharides yeast extract from Millipore. This suggests a higher β-glucan content in the Millipore sample compared to the other samples, likely caused by indiscriminate polysaccharide precipitation. Supporting this hypothesis, the mannose-to-glucose ratio for the same yeast extract processed via UF was substantially higher (about 10-fold higher). Notably, the broad peak at the lower Mw end (∼39 min) in the SEC profile of TCA/EtOH extracted polymers (**Figure 1B**)is absent in the UF-extracted polymers (**Figure 1A**). This is most likely β-glucan that was largely removed by the 30 kDa ultrafiltration membrane.

### 3.3 ^1^H-NMR spectroscopic analysis verifies the presence of high Mw α-mannan and β-glucan in the commercial yeast extract

To support identification of the polysaccharides detected by SEC (**Figure 1**) that were partially characterized by monosaccharide composition analysis (**Figure 2** and **Table 2**), ^1^H-NMR spectroscopic analysis was carried out for all samples. The polysaccharides derived from yeast extracts revealed peak patterns closely resembling those of commercial yeast mannan, regardless of the extraction method and supplier (**Figure 3**). UF-extracted polymers also contain minor amounts of β-glucan (**Figure 3B, D, E**), as indicated by the doublet at 4.54 ppm. This doublet was larger in the polymer mixture extracted by the TCA/EtOH method (**Figure 3C)**, suggesting a higher amount of β-glucan, aligning well with the observations made by SEC and HPAEC-PAD. Additionally, the yeast extract retentate from Oxoid™ displayed signal characteristics corresponding to amino acid side chains of high Mw peptides (**Figure 3E**), which aligns with SEC data showing peaks eluting at ∼37–40 min with high absorbance at 280 nm (**Figure 1D**).

**Figure 3.**
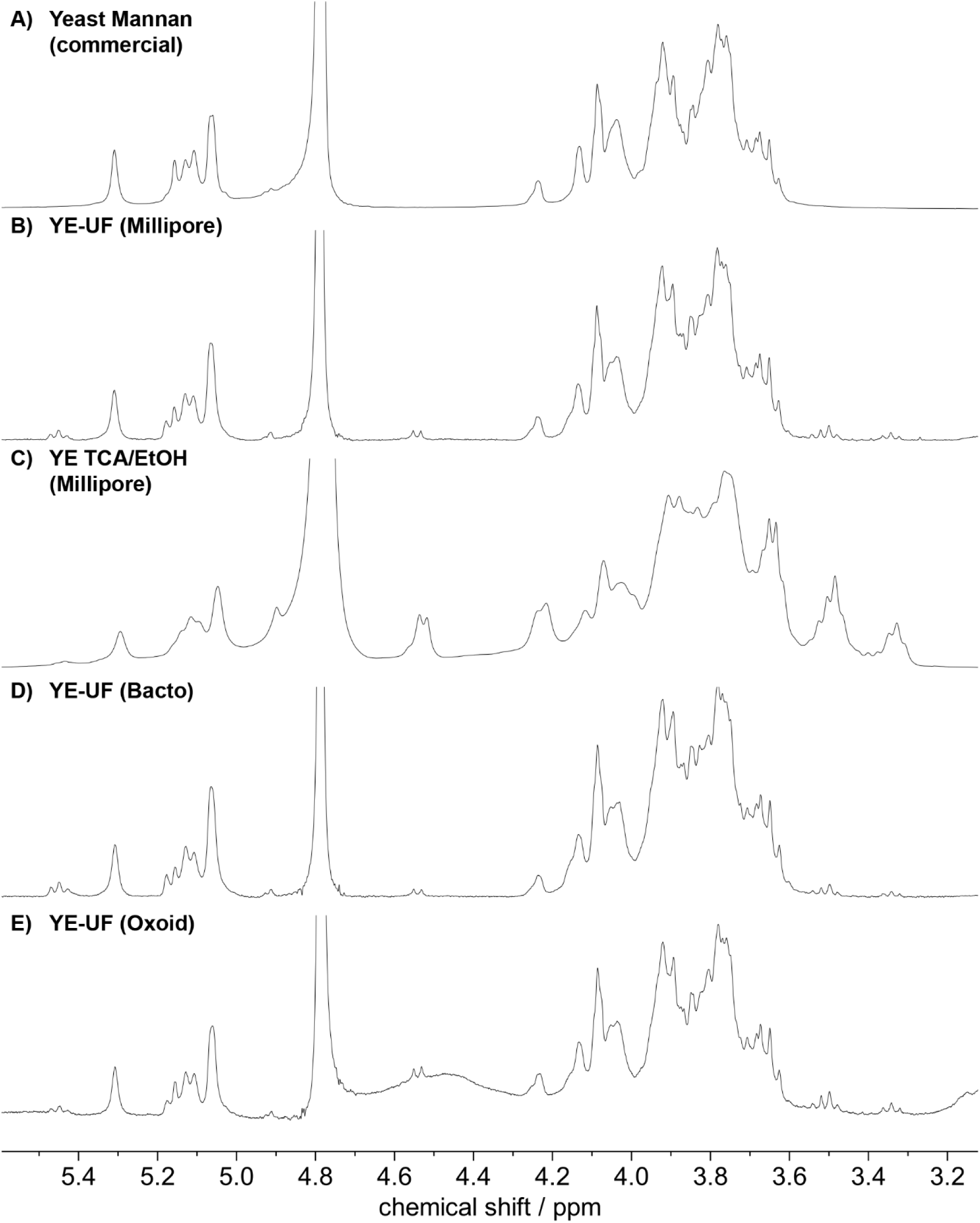
^1^H-NMR spectra (D_2_O, 400 MHz, 298 K) of commercial yeast mannan (**A**), recovered polymers from various yeast extracts, obtained using ultrafiltration (**B**, **D**, and **E**) and TCA/EtOH method (**C**). YE, yeast extract; UF, ultrafiltration; MRS, De Man, Rogosa and Sharpe medium.

Yeast mannan from *Saccharomyces cerevisiae*, which is composed of a 1,6 linked α-mannan backbone with a high degree of α-1,2 and α-1,3 mannan branches, can be easily identified in an ^1^H-NMR-spectrum by using the anomeric signals as a fingerprint (**Figure 4**). The pattern to look out for when identifying yeast mannan is an arrangement of four smaller peaks between 5.10 and 5.20 ppm which is flanked by two larger peaks at 5.06 and 5.31 ppm. At 4.91 ppm, a signal stemming from 1,6 linked unbranched mannose is also observed, which is usually small due to the highly branched nature of the polysaccharide. Native yeast mannan is also phosphorylated to a low degree which is indicated by the signals at 5.45 ppm. This phosphorylation is often absent in commercial yeast mannan, as these products are usually prepared by alkaline extraction, which leads to dephosphorylation.

**Figure 4.**
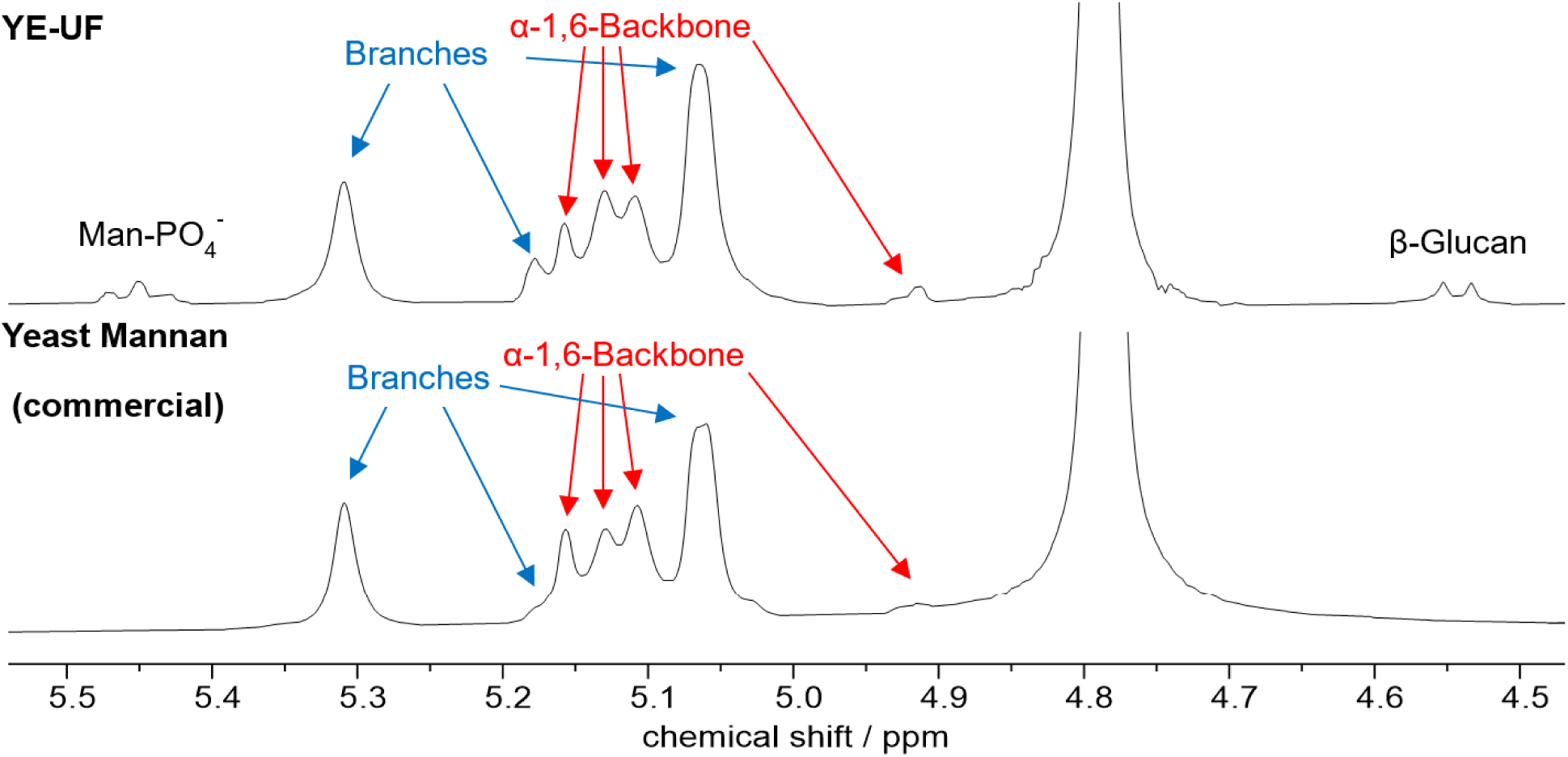
Anomeric signals in ^1^H-NMR (D_2_O, 400 MHz, 298 K) of high Mw polymer in yeast extract (top) and commercial yeast mannan (bottom). Signals from the backbone mannoses are marked in red while signals from branches are marked in blue. Annotations are based on a study by Vinogradov *et al.* [47].

Since the absolute chemical shift depends on the calibration, it is important to compare the relative difference in chemical shift between peaks when comparing shift values to identify yeast mannan (**Figure 5**). Minor differences in relative peak intensities may also be observed due to slight differences in branching.

**Figure 5.**
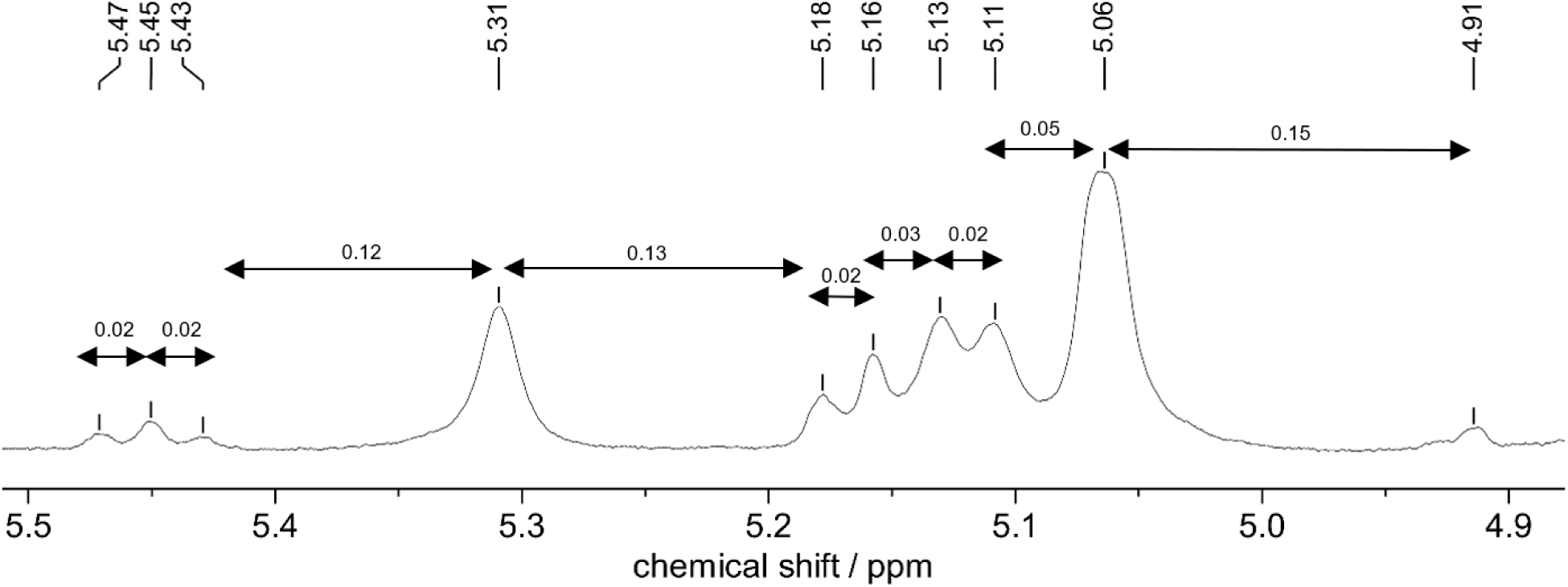
Anomeric signals in ^1^H-NMR (D_2_O, 400 MHz, 298 K) of native yeast mannan with relative differences in chemical shift annotated in ppm.

Some publications reporting microbial EPS structures present ^1^H-NMR spectra that exclusively show signals corresponding to yeast mannan [3,6,9,13,35,42,52]. Others display ^1^H-NMR spectra containing signals characteristic of yeast mannan in conjunction with other signals, suggesting that these EPS extracts are contaminated [8,12,18,30,36,48,51]. There are also publications of bacteria that seemingly produce mannan [23,46]. Based on SEC data reported for EPS in these publications (**Table 1**), yeast mannan was identified across peaks corresponding to a broad range of molecular weights. These findings are not entirely consistent with one another and also differ from our observations in SEC profiling. Based on our findings (**Figure 1**), if the EPS produced has a Mw >500 kDa, it will most likely not include yeast mannan contaminant, as these are lower in Mw (∼50–100 kDa). However, it would require further purifications using ultrafiltration, dialysis, or SEC fractionation in order to separate the EPS from the contaminant.

### 3.4 ^1^H-NMR spectroscopic analysis of EPS extract can reveal α-mannan contamination

An example of how severely yeast mannan can interfere with the interpretation of structural data is shown by the EPS extract from *L. rhamnosus* that was cultivated in conventional MRS medium (**Figure 6**). The ^1^H-NMR spectrum of the resulting “EPS extract’’ showed yeast mannan signals overlapping with actual EPS signals (**Figure 6, middle**).

**Figure 6.**
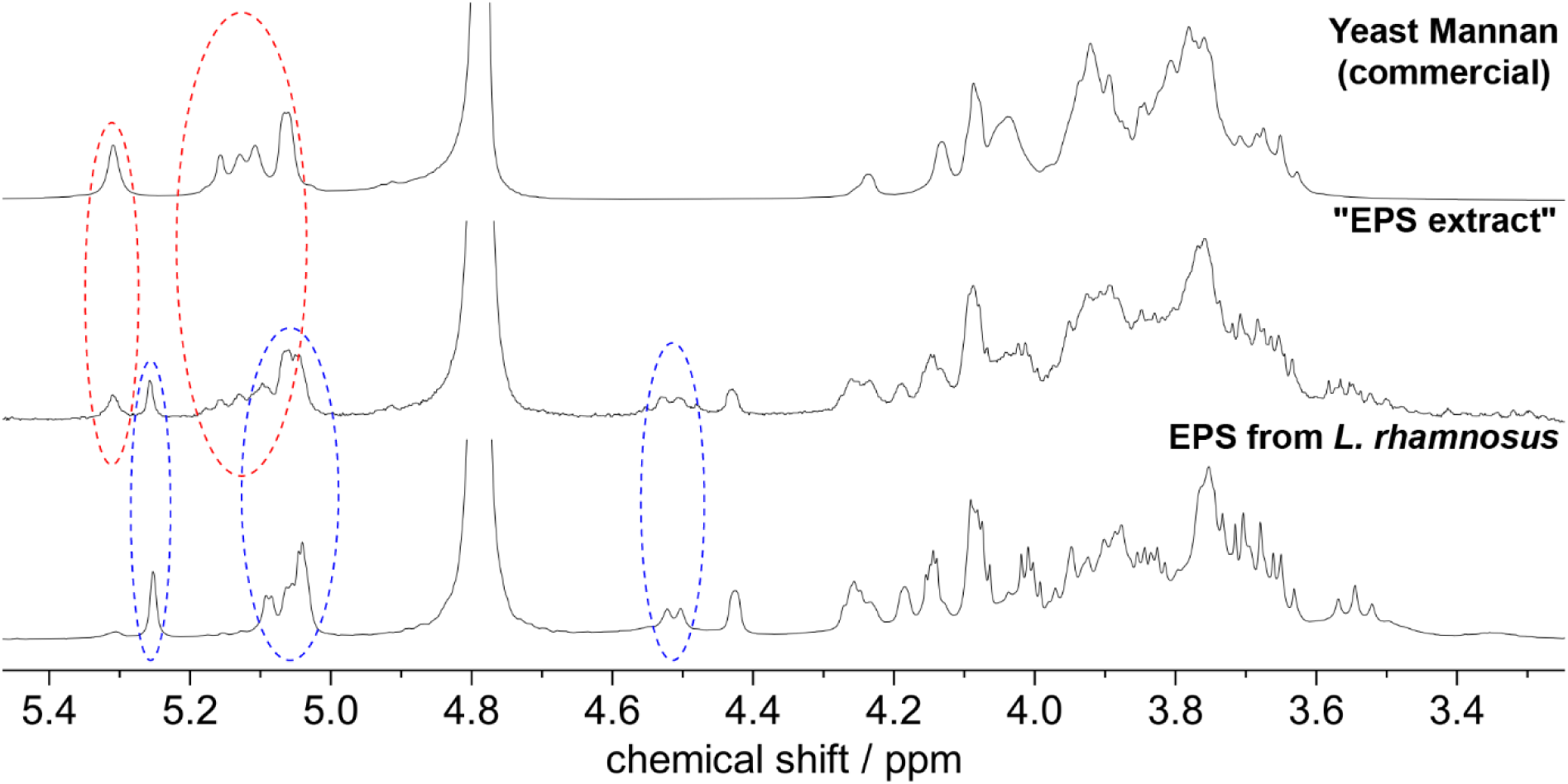
^1^H-NMR (D_2_O, 400 MHz, 298 K) of commercial yeast mannan (top), EPS extract from *L. rhamnosus* that was cultivated in conventional MRS medium (middle), and purified EPS from *L. rhamnosus* (bottom). Anomeric signals from yeast mannan are highlighted in red, while anomeric signals from the actual EPS are highlighted in blue.

The presence of yeast mannan was easily identified by peaks in the anomeric region (**Figure 6, highlighted in red**), which partially overlapped with anomeric peaks from the actual EPS (**Figure 6, highlighted in blue**) in this case. Fractionation of the high Mw peak (**Figure 1E**) using preparative SEC yielded pure exopolysaccharide from *L. rhamnosus*, whose ^1^H-NMR spectrum (**Figure 6, bottom**) matched the previously published spectrum [24]. However, purification by SEC is only feasible if the EPS of interest has a Mw that differs significantly from that of yeast mannan. For isolation of EPS with similar molecular weight, removal of the high Mw contaminants from the medium prior to fermentation may be necessary (see Section 3.5).

### 3.5 UF-MRS is a superior option as growth medium for obtaining LAB-EPS of high purity

In line with other studies [5,31], ultrafiltration was employed to remove high Mw polysaccharides from the yeast extract prior to bacterial cultivation. In our case, the UF-MRS medium was prepared by ultrafiltration of the conventional MRS medium with a 30 kDa Mw cut-off membrane. The resulting permeate and retentate were analyzed by SEC to estimate the molecular weight distribution of their components. SEC analysis qualitatively indicated that most compounds in the permeate had molecular weights around 1.3 kDa (**Figure 7**).

**Figure 7.**
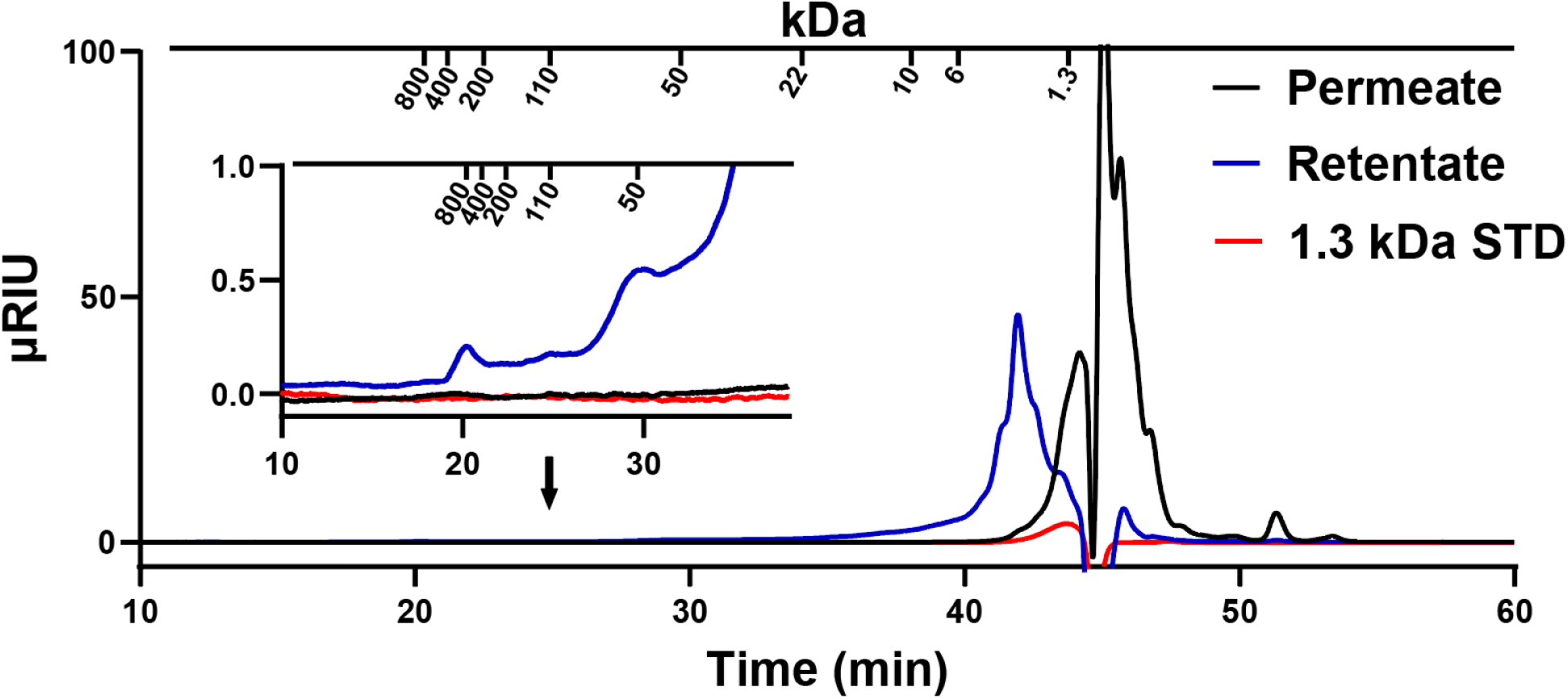
SEC overlay of permeate (black trace) and retentate (blue trace) from MRS medium after ultrafiltration, along with 1.3 kDa pullulan standard. The zoom in view of chromatogram displays the presence of high Mw contaminants in the retentate. Separation was performed by Yarra 4000-2000 series (300 × 7.8 mm, 3 µm) using 0.1 M Na_2_SO_4_ as mobile phase. Peak elution time of pullulan standards (Mw in kDa) are marked above chromatogram; from left to right 800, 400, 200, 110, 50, 22, 10, 6 and 1.3kDa, respectively.

The permeate (UF-MRS) was subsequently used for the cultivation of *L*. *pentosus* KW1, resulting in growth curves comparable to that observed with untreated MRS medium. The same ultrafiltration membrane (30 kDa Mw cut-off) was also used during EPS purification, where the retentate was collected following diafiltration. For *L. pentosus*, three distinct peaks were fractionated by preparative SEC (**Figure 1F–G**) and further characterized using NMR (**Figure 8**). Without an ultrafiltration step of the MRS medium prior to fermentation, **peak 2** (∼50 kDa) contained mannan (**Figure 8, middle**). Ultrafiltration of the medium prior to cultivation proved effective in removing high Mw contaminants that would otherwise compromise the purity and structural characterization of the produced polymer (**Figure 8, bottom**).

**Figure 8.**
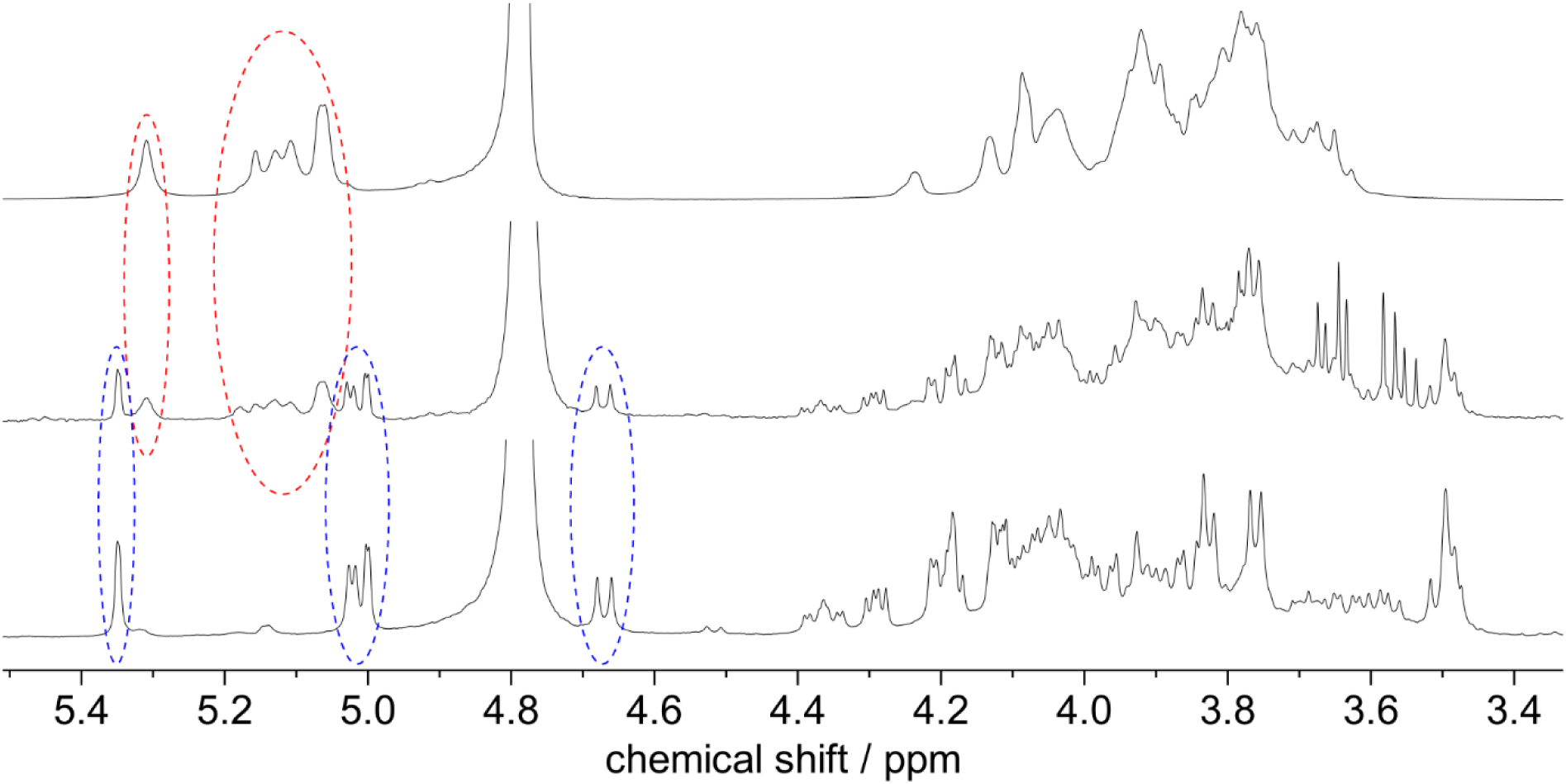
^1^H-NMR (D_2_O, 400 MHz, 298 K) of commercial yeast mannan (top), peak 2 of *L. pentosus* cultivated in conventional MRS medium (middle), and peak 2 of *L. pentosus* cultivated in UF-MRS (bottom). Anomeric signals from yeast mannan are highlighted in red, while anomeric signals from the actual polymer are highlighted in blue.

The ultrafiltration step may not be necessary if the bacterial strain is capable of utilizing mannan contaminants as a carbon source; however, this would require dedicated characterization of the strain prior to EPS production. Alternatively, chemically defined ingredients, such as yeast nitrogen base, can be used for EPS production [20,40]. Regardless of the medium used, it is always advisable to include a non-inoculated medium as a control to account for background components and ensure accurate interpretation of results.

In this study, we demonstrate that yeast extracts commonly used in bacteriological media contain high Mw polysaccharides, primarily mannans, which can contaminate EPS preparations. If not metabolized by the bacteria, these contaminants remain in the fermentation broth and may be co-purified with the EPS, potentially leading to compromised purity, misinterpretation of structural data, and potentially misleading results if used in applications or experiments. We show how ^1^H-NMR can be used to validate the presence of yeast α-mannans in the purified product. We also demonstrate that ultrafiltration of the culture medium prior to bacterial cultivation is a convenient and efficient way to prevent high Mw contaminants in purified EPS.

## Acknowledgement

The authors thank Dr. Kamilla Wiull and Prof. Geir Mathiesen for kindly providing bacterial stocks, and Synnøve Sætervik and Cornelia Rognstad Karlsen for their involvement in EPS production.

