## Supplementary file for "High molecular-weight polysaccharide contamination from yeast extract in semi-defined bacteriological media: Effects on exopolysaccharide production and purity"

### Supplementary Materials

**Table S1.** An extended overview of studies reporting microbial EPS that may be contaminated by mannan. Includes reported NMR linkages and suggested biological/physical effects. Lactobacillus studies (gray highlight), Gram-positive bacteria (green highlight), Gram-negative bacteria and fungi (blue highlight). Abbreviations: YM, yeast mannan; YE, yeast extract; WP, whey permeate; LB, Luria-Bertani broth; NB, nutrient broth.

| Species/Strain | Cultivation Media | Reported linkages[1] | Suggested biological/physical effects | Ref. |
|---|---|---|---|---|
| *L. plantarum* VAL6 | MRS | β-D-Glc and β-D-Gal*p*. No linkages reported. | Antioxidant activity | [34] |
| *L. plantarum* 70810 | YE based medium | EPS1: 1,6-linked-α-D-Glc*p*, 1,2-linked-α-D-Man*p*, 1,3-linked-α-D-Man*p*, 1,4-linked-α-D-Gal*p*, 1,3,4-linked-α-D-Man*p*, 1,6-linked-α-D-Man*p* and terminal 1-linked α-D-Man*p*<br>EPS2: 1,6-linked-α-D-Glc*p*, 1,4-linked-β-D-Gal*p*, 1,3,6-linked-α-D-Gal*p*, 1,6-linked-α-D-Gal and 1,3,4-linked-α-D-Gal*p* and terminal α-D-Man*p*-1 | Antioxidant and antitumor activity | [48] |
| *L. plantarum* BC-25 | MRS | 1,2-linked-β-D-Man*p*, 1,2-linked-α-D-Glc*p*, 2,6-linked-β-D-Man*p* and 2,6-linked-α-D-Gal*p*. Note: 2-6 linkages in polysaccharides are very unlikely. | Selenium uptake for bioavailability | [53] |
| *L. plantarum* HMX2 | MRS | No data. | - | [50] |
| *L. plantarum* MTCC 9510 | YE based medium | 1,3-linked-α-D-Man*p*, 1,3-linked-α-D-Glc*p* and 1,3-linked-β-D-Glc*p* | - | [19] |
| *L. plantarum* W1 | MRS + YE | 1,2,6-linked-α-D-Glc*p*, 1,3-linked-α-D-Man*p*, 1,3-linked-α-D-Glc*p*, 1,6-linked-α-D-Man*p* and terminal 1-linked-α-D-Glc*p* | - | [12] |
| *L. helveticus* MB2-1 | WP + YE | No data. | Antioxidant and metal ion chelating activity | [29] |

| | | | | |
|---|---|---|---|---|
| *L. paracasei* 2333<br>*L. rhamnosus* 1019<br>*L. bulgaricus* 1932 | Modified MRS | No data. | - | [15] |
| *L. rhamnosus* ACS5 | MRS | 1,4-linked-α-D-Man*p*, 1,4-linked- α-D-Man*p*NAc and 1,6-linked-α-D-Gal*p* | Antidiabetic and antioxidant activity | [18] |
| *L. paracasei* EPS DA-BACS | MRS | 1,2-linked-α-D-Man*p*, 1,6-linked-α-D-Man*p*, 1,2,6-linked-α-D-Man*p* and terminal 1-linked-α-D-Glc*p* | Anti-inflammatory and antimicrobial activities | [26] |
| *L. crispatus* L1 | YE based medium | 1,6-linked-α-D-Man*p*, 1,2-linked-α-D-Man*p*, 1,3-linked-α-D-Man*p,* 1,2,6-linked-Man*p* and terminal 1-linked-α-D-Man*p* | Probiotic | [13] |
| *Lactobacillus spp.* | MRS | 1,3-linked-α-D-Man , terminal 1-linked-β-D-Man and terminal 2-linked-α-D-Glc<br>Note: The proposed structure is a trisaccharide, not a polymer. | Anti-tumor | [44] |
| *Lactobacilli strains* | MRS | No data. | Anti-biofilm | [41] |
| *S. thermophilus* ASCC 1275 | M17 | Fraction 1: 1,2-linked-α-D-Man*p*, 1,2,6-α-D-linked-Man*p* and terminal 1-linked-α-D-Man*p*<br>Fraction 2: 1,2,6-α-D-linked-Man*p* and terminal 1-linked-α-D-Man*p*<br>Fraction 3: 1,3,6-linked-β-D-Glc*p*, 1,3-linked-β-D-Gal*f,* 1,3-linked-α-D-Glc*p,* 1,6-linked-β-D-Glc*p,* 1,4,linked-β-D-Gal*p* and terminal 1-linked-β-D-Gal*p* | - | [36] |
| *E. faecium* MS79 | MRS | 1,4-linked-α-D-Glc, 1,3-linked-α-D-Glc, 1,2,4-linked-β-D-Man, 1,2-linked-α-D-Glc, 1,6-linked-α-D-Glc and terminal 1-linked-α-D-Gal | Anti-cancer and anti-diabetic properties | [1] |
| *E. faecalis* | LB | 1,4-linked-D-Man, 1,6-linked-D-Man, 1,3,4-linked-D-Man, terminal 1-linked-D-Glc and terminal 1-linked-L-Fuc. | Antioxidant, metal ion chelation, and hydroxyl radical scavenging activity | [8] |

| | | | | |
|---|---|---|---|---|
| *B. breve* H4–2 | MRS | 1,2-linked-α-D-Man*p*, 1,3-linked-α-D-Man*p*, 1,2,6-linked-α-D-Man*p*, 1,6-linked-α-D-Glc*p* and terminal 1-linked-α-D-Man*p* | Potential immune-stimulating polysaccharide | [35] |
| *P. acidilactici MT41-11* | MRS | EPS-1: 1,3-linked-α-D-Man*p*, 1,2-linked-α-D-Man*p* and terminal 1-linked-α-D-Man*p*<br>EPS-2: 1,3-linked-α-D-Glc*p*, 1,2-linked-α-D-Glc*p*, terminal 1-linked-α-D-Glc*p*, 1,2-linked-α-L-Fuc*p*, 1,6-linked-β-D-Glc*p* and terminal 1-linked-β-D-Glc*p* | Prebiotic, antioxidant, anti-biofilm | [3] |
| *B. licheniformis* | MRS | No data. | - | [25] |
| *Tetragenococcus halophilus* | MRS | EPS-1: Terminal 1-linked-α-D-Glc*p* 1,4-linked-β-D-Glc*p,* 1,4,6-linked-α-D-Glc*p*, 1,3,6-linked-β-D-Glc*p*, 1,6-linked-α-D-Glc*p*, 1,2-linked-α-D-Man*p*, 1,2,6-linked-α-D-Man*p*, 1,6-linked-α-D-Man*p* and terminal 1-linked-α-D-Man*p*<br>EPS-2: 1,2-linked-α-D-Man*p*, 1,2,6-linked-α-D-Man*p*, 1,3,6-linked-α-D-Man*p*, 1,6-linked-α-D-Man*p*, terminal 1-linked-α-D-Man*p* and terminal 1-linked-β-D-Man*p* | Antioxidant activity | [51] |
| *Rhodopseudomonas palustris* | Salts + YE | 1,2-linked-Man*p*, 1,4-linked-Man*p*, 1,6-linked-Man*p*, 1,2,6-linked-Man*p*, and terminal 1-linked-Man*p* | Immunomodulator and prebiotic | [52] |
| *P. haloplanktis* TAC 125 | YE based medium | 1,6-linked-α-D-Man*p*, 1,2-linked-α-D-Man*p*, 1,3-linked-α-D-Man*p*, 2,6-linked-α-D-Man*p*, terminal-D-Man*p*, terminal 1-linked-β-D-Glc*p* and 1-P-D-Man*p* | - | [9] |
| *Psychrobacter arcticus* 273-4 | LB | 1,6-linked-α-D-Man*p*, 1,2-linked-α-D-Man*p*, 1,3-linked-α-D-Man*p*, 2,6-linked-α-D-Man*p*, terminal-D-Man*p*, terminal 1-linked-β-D-Glc*p* and 1-P-D-Man*p* | Ice recrystallization inhibition activity | [6] |
| *Shigella flexneri* CMCC51574 | LB | terminal 1-linked-α-D-Glc*p*, terminal 1-linked-α-D-Man*p*, 1,2,6-linked-α-D-Man*p*, 1,3-linked-α-D-Man*p* and 1,2-linked-Man*f*. Note: The reported $^{13}$C-NMR-shifts of C1 and C5 are way too low for a furanose. | Biofilm formation | [42] |

| | | | | |
|---|---|---|---|---|
| *A. sinodeliciosus* var. Chaidam | YE based medium | Main chain: 1,2,6-linked-α-D-Man*p*, 1,6-linked-α-D-Gal*p*, 1,2-linked-α-D-Gal*p*, and 1,6-linked-α-D-Man*p* 1,2,6-linked-α-D-Gal*p*, terminal 1-linked-α-D-Man*p*, terminal 1-linked-α-Gal*p*, and 1,3-linked-β-Gal*p*NAc. | Neuroprotective activity | [30] |
| *P. aeruginosa* AG01 LC586427 | NB | No data. | Antibacterial, anticancer, and antiviral activities | [16] |

[1] The various exopolysaccharides mentioned in the literature, found in this table, use various abbreviations, such as EPS 1, 2 etc. The numbering typically follows the chronological elution order based on SEC, with EPS 1/Fraction 1 representing the fraction with the highest Mw.
